# Comprehensive single-cell atlas of the mouse retina

**DOI:** 10.1101/2024.01.24.577060

**Authors:** Jin Li, Jongsu Choi, Xuesen Cheng, Justin Ma, Shahil Pema, Joshua R. Sanes, Graeme Mardon, Benjamin J. Frankfort, Nicholas M. Tran, Yumei Li, Rui Chen

**Author notes:** These authors contributed equally to this work (Jin Li, Jongsu Choi, Xuesen Cheng). Corresponding author: Rui Chen.

## Abstract

Single-cell RNA sequencing (scRNA-seq) has advanced our understanding of cellular heterogeneity at the single-cell resolution by classifying and characterizing cell types in multiple tissues and species. While several mouse retinal scRNA-seq reference datasets have been published, each dataset either has a relatively small number of cells or is focused on specific cell classes, and thus is suboptimal for assessing gene expression patterns across all retina types at the same time. To establish a unified and comprehensive reference for the mouse retina, we first generated the largest retinal scRNA-seq dataset to date, comprising approximately 190,000 single cells from C57BL/6J mouse whole retinas. This dataset was generated through the targeted enrichment of rare population cells via antibody-based magnetic cell sorting. By integrating this new dataset with public datasets, we conducted an integrated analysis to construct the Mouse Retina Cell Atlas (MRCA) for wild-type mice, which encompasses over 330,000 single cells. The MRCA characterizes 12 major classes and 138 cell types. It captured consensus cell type characterization from public datasets and identified additional new cell types. To facilitate the public use of the MRCA, we have deposited it in CELLxGENE, UCSC Cell Browser, and the Broad Single Cell Portal for visualization and gene expression exploration. The comprehensive MRCA serves as an easy-to-use, one-stop data resource for the mouse retina communities.

## Introduction

The retina is a highly heterogenous part of the eye that captures and processes the light signal ^1–3^. The processing is enabled through five classes of retinal neurons: photoreceptors (PR), horizonal cells (HC), bipolar cells (BC), amacrine cells (AC), and retinal ganglion cells (RGC), which form an intricate circuitry necessary for processing and relaying the light signal to the visual cortex. Non-neuronal cells such as Müller glia cells (MG), microglia, astrocytes, and retinal pigment epithelial cells (RPE) provide structural integrity of the tissue and carry out various supporting roles such as metabolism and neuronal homeostasis in the retinal microenvironment ^4,5^. Characterization of distinct retinal cell types is, therefore, critical in advancing our understanding of the fine intricacies of cell interactions involved in retinal biology and visual disorders.

Single cell technologies have opened a window into knowledge of cellular heterogeneity and intricate cell-to-cell interactions that cannot currently be resolved at the tissue level and have allowed exploration of individual cellular expression signatures, which can be mapped to unique molecular cell types ^6,7^. The resulting cell atlas can serve as a foundation for numerous applications, including the annotation of cell types in other scRNA-seq experiments ^8^, the identification of differentially expressed targets for purification or manipulation ^9^, and the generation of marker panels useful for single-molecule imaging, including spatial profiling ^10^. While studies have demonstrated cell type heterogeneities in various tissues, several perplexing issues remain to be addressed in establishing a comprehensive cell atlas such as the agreement on cell type definitions across different experiments or whether enough cells have been profiled to exhaust all existing cell types. Integrated analyses of various scRNA-seq datasets from different studies, therefore, can provide an important insight that comprehensively addresses such issues.

The mouse retina provides an important model for the study of neurobiology, with more than 130 distinct cell types characterized through previous scRNA-seq studies ^7,9,11–15^. However, the scRNA-seq datasets have been generated separately for BC ^11^, AC ^12^, and RGC ^9,13,14^, with the largest dataset containing just under 36,000 cells, making it difficult to use in aggregate. Though most of these datasets are independently browsable on the Broad Single Cell Portal ^16^ and accessible through separate databases such as the Gene Expression Omnibus (GEO) repository, it can be challenging to assess gene expression patterns across all retinal cell types. Ensuring these atlases define a complete set of retinal cell types remains a major challenge that can only be addressed by powering studies to sufficiently profile the rarest retinal cell types. Here, we generated scRNA-seq data of over 189,000 cells in the mouse retina to complement 141,000 cells from six publicly available scRNA-seq datasets ^9,11–15^, creating a unified cell atlas of the wild-type mouse retina containing over 330,000 cells. Our integrated analysis presents a comprehensive characterization of all major cell classes in the retina, including non-neuronal types, as well as a consensus cell type annotation of BCs, ACs, and RGCs. Accessible, interactive web browsers have facilitated easy visualization of atlas characterizations and exploration of gene expression in the MRCA. The comprehensive unified MRCA will serve as a valuable resource for the community.

## Results

### Generation of scRNA-seq dataset for wild-type mouse retina

To establish a comprehensive atlas of the mouse retina, we performed scRNA-seq profiling with C57BL/6J mouse retina tissue samples, aged from P14 to 12 months, for over 189,000 cells (**Fig. 1a** and Methods). As summarized in **Table 1**, six samples of varying ages were dissociated retinal cells without enrichment, and ten samples of eight weeks old were enriched using surface markers CD73 and CD90.1 to enrich for rare cell population. Depletion of rod photoreceptors was achieved by removing cells positive for CD73 using anti-CD73-PE antibody and anti-PE magnetic beads, which primarily label photoreceptor precursors and mature rod photoreceptors in mice ^12,17^. To enrich ACs and RGCs, CD90.1 positive cells are selected ^18,19^.

**Figure 1.**
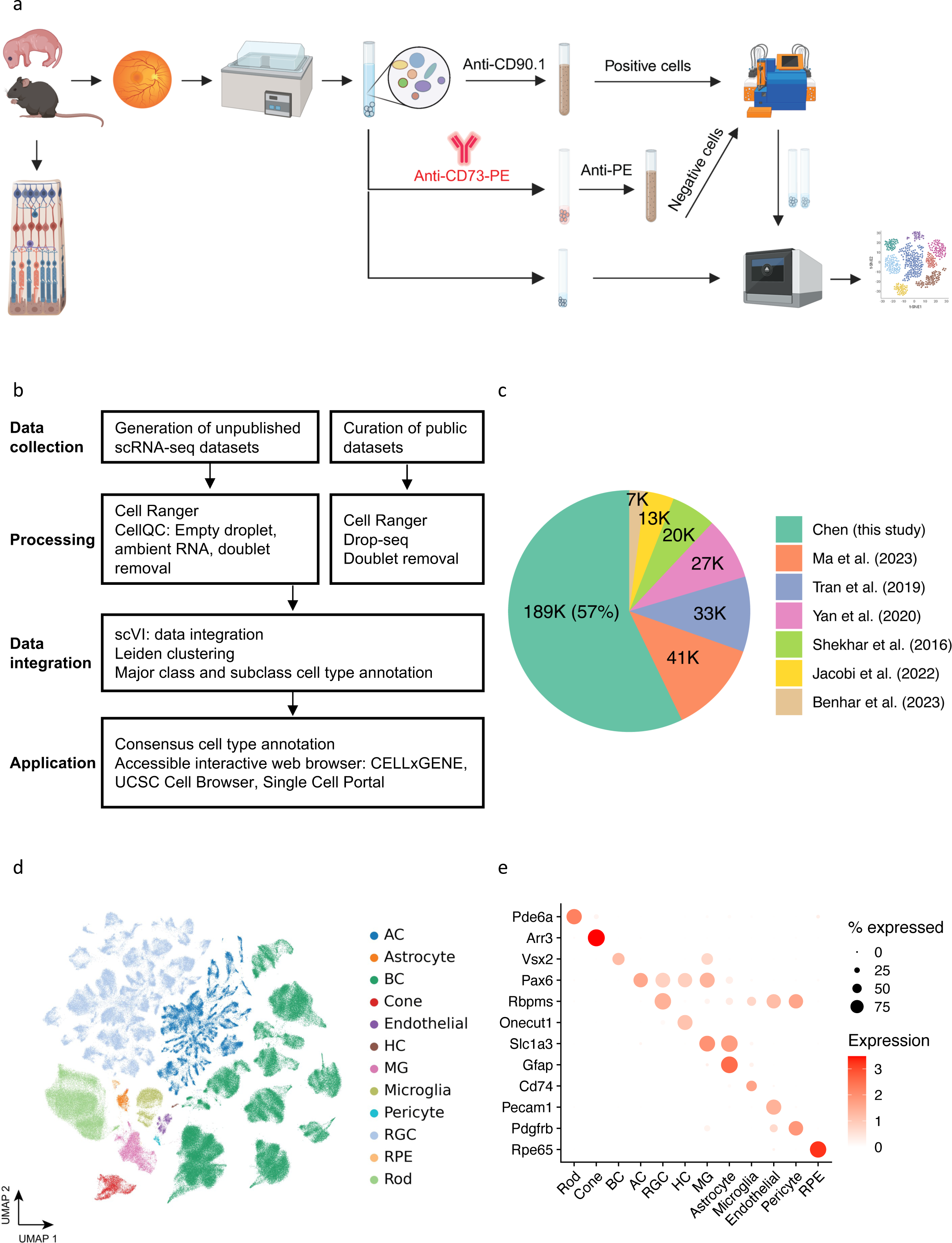
Overview of single cell atlas of the mouse retina. **(a)** The workflow for generating unpublished scRNA-seq datasets. The data generation process involved using mice aged from P14 to 12 months. Following retina dissection and cell dissociation, single cells were enriched using autoMACS with Anti-CE73-PE antibodies or Anti-CD90.1 beads for specific amacrine, bipolar, and retinal ganglion cells. Subsequently, 10X single-cell RNA sequencing was performed on both the unenriched and enriched single cells. The retained single cells were then utilized in downstream atlas construction. **(b)** The integrated analysis workflow for constructing the MRCA. To construct a comprehensive unified single-cell reference of the mouse retina, we generated 16 unpublished scRNA-seq samples of the mouse retina and incorporated four curated public datasets to enhance specific amacrine, bipolar and retinal ganglion cells. The collected data were processed using the Cell Ranger and CellQC pipeline to produce feature count matrices. Feature counts were then processed to remove estimated empty droplets, ambient RNA, and doublets. The retained cells were integrated using scVI to eliminate batch effects across samples. The trained low-dimensional embeddings were used to calculate cell dissimilarities and identify clustering through a two-level clustering approach. Major class and subclass cell types were annotated using canonical marker genes and public labeling. To facilitate user-friendly access and exploration, the MRCA was deployed on accessible interactive web browsers, including CELLxGENE, UCSC Cell Browser, and Single Cell Portal. **(c)** Pie chart displaying the percentage of cells contributed by each dataset used in the MRCA. **(d)** UMAP visualization of the MRCA colored by major classes. **(e)** Dot plot illustrating the expression of canonical markers for major classes.

### Integration of scRNA-seq datasets for the mouse retina

To compile the most comprehensive scRNA-seq data for the MRCA, we curated and obtained six publicly available scRNA-seq datasets, each enriched for a specific cell type using transgenic labels or immunolabeling combined with FACS. Together, they consisted of over 141,000 cells. To consolidate the transcript annotation between different datasets, we used the Cell Ranger (version 7.0.1) pipeline to align raw FASTQ files from four datasets obtained from GEO and Sequence Read Archive (SRA) repositories. Count matrices of these datasets were generated using the mm10 reference genome obtained from 10x Genomics (https://cf.10xgenomics.com/supp/cell-exp/refdata-gex-mm10-2020-A.tar.gz). Five of the published studies were sequenced on the 10x Genomics 3’ platform, and one (Shekhar *et al.*) was generated using the Drop-seq protocol ^7^. The Drop-seq data were aligned against mm10 and processed into count matrices using the Drop-seq pipeline (https://github.com/broadinstitute/Drop-seq). The cell type labels of previous annotations were obtained from the Broad Single Cell Portal website ^16^. To remove technical variations introduced across different experiments or studies, scVI ^20^ was applied to integrate all newly generated and public datasets, generating a low-dimensional representation (**Fig. 1b** and Methods). Putative cell doublets were further removed using the deep learning doublet identification method Solo ^21^ (**Supplementary Fig. 1a**).

In the integrated data, the public dataset accounts for 43%, while the newly generated data accounts for the remaining 57% (**Fig. 1c**). Within the integrated UMAP, 97 clusters were identified (**Supplementary Fig. 1b**). These clusters were annotated as one of 12 major classes, including PR, BC, AC, RGC, HC, MG, RPE, astrocyte, microglial, endothelial, and pericyte, using known marker gene expression ^22,23^ (**Supplementary Fig. 1c**). Cells from non-enriched retina samples showed a distribution across major classes at an expected proportion, with rod photoreceptors as the biggest proportion ^2^. In contrast, enriched samples from both newly generated data and previous studies showed the expected skewed distribution of cell types in BCs, ACs, and RGCs (**Supplementary Fig. 1d**). The two newly generated samples with enrichment methods, CD73^-^ and CD90.1^+^ samples, were primarily composed of BCs and ACs, respectively, contributing to 83% (122.6K out of 147.7K) and 25% (11.2K out of 44K) of all BCs and ACs in the integrated data, respectively.

Previous studies have identified 15 distinct types of BCs, 64 ACs, and 46 RGCs ^9,11,12^. To determine the consensus annotation of neuronal types for these subclasses, we performed clustering analysis at higher resolution within individual BC, AC, and RGC classes (**Fig. 1d** and **Fig. 1e**).

### 15types of bipolar cells

A total of 147,700 BCs were identified in the integrated datasets, with 122,600 cells from our newly generated CD73^-^ sample and 19,800 cells from the Shekhar *et al.* study ^11^. The integrated analysis identified 15 BC clusters, corresponding to previously annotated BC types (**Fig. 2a-b** and Methods). The 15 clusters of integrated BCs showed a generally even distribution of cells from various samples, with the exception of two types, BC1A and BC1B, where more than 90% of populations came from the study by Shekhar *et al.* possibly due to differences in enrichment methods (**Fig. 2a, 2d** and **Supplementary Fig. 2d-e**). The final annotation of BCs revealed consistent expression profiles of previously identified BC type marker genes ^11,24^ (**Fig. 2b-c**). With a significant addition of BCs in the MRCA, clear separation of BC8 and BC9 is observed, which were merged but demonstrated substructure in the Shekhar et al. dataset (**Fig. 2a-b**). The separate clusters showed proper expression patterns of known markers like *Cpen9* in BC9 ^11,25^. In addition, additional BC type markers were identified via differential gene expression analysis, which showed more specific expressions than previous marker genes, such as *Tafa4* in BC4, *Ptprt* in BC5A, and *Gm13986* in BC8 (**Fig. 2e**). Interestingly, despite an almost ten-fold increase in the number of BCs in our analysis, we did not observe any sign of a novel cell type, which suggests that the mature mouse retina likely only contains 15 BC types.

**Figure 2.**
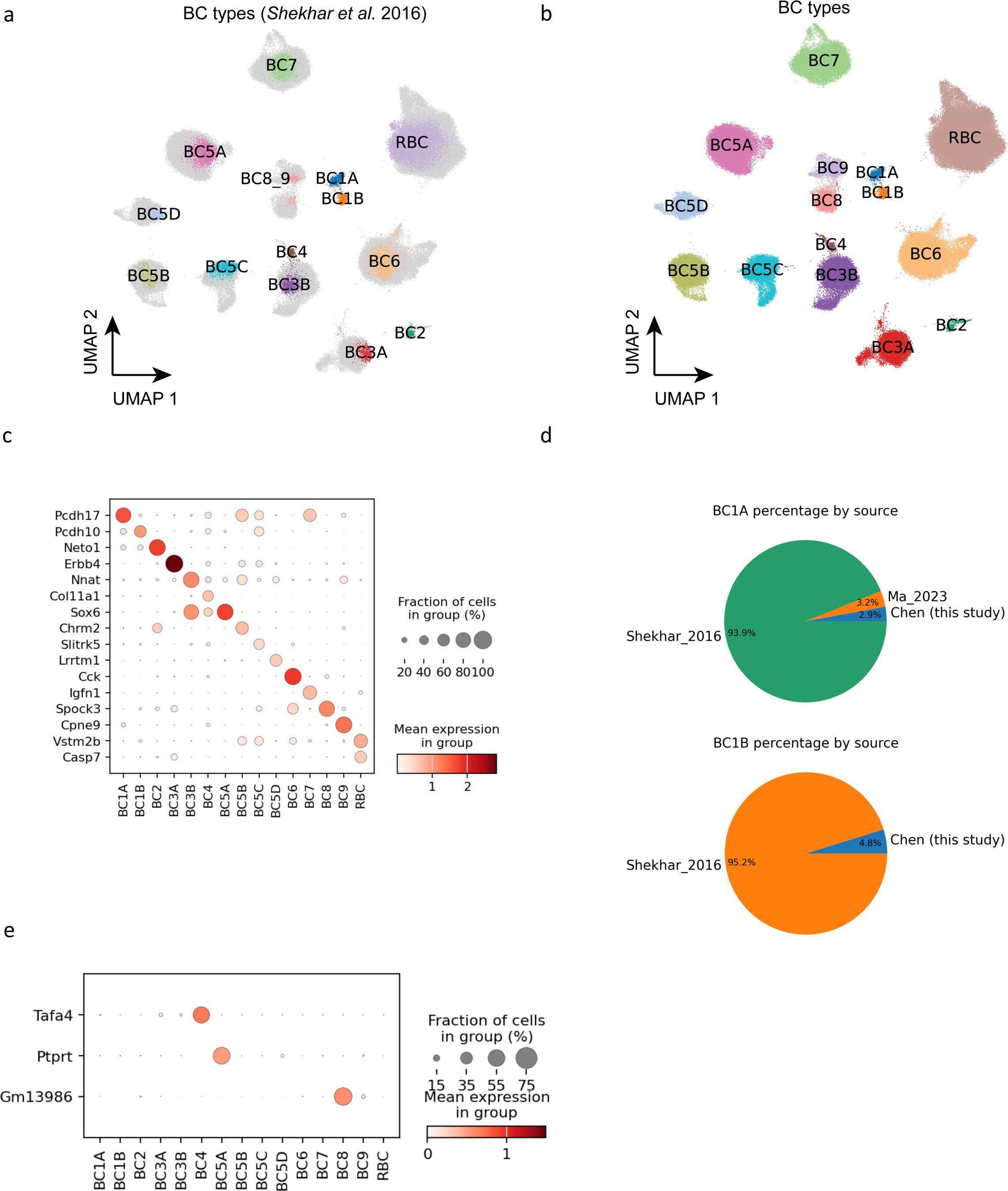
Bipolar cells. **(a)** UMAP visualization of BCs colored by public cell type labels from *Shekhar et al.* 2016. The newly discovered cells without public labeling are colored in gray. **(b)** BCs colored by the 15 annotated annotated cell types. **(c)** Dot plot of BC type marker gene expression in the 15 types. **(d)** Pie chart showing the percentage each data source making up BC1A and BC1B population. **(e)** Dot plot of new markers for three BC types: BC4, BC5A, and BC8. The three new markers exhibit more exclusive expression patterns.

### Amacrine cells

Through CD90.1 positive enrichment, the newly generated samples contributed 11,200 ACs, in addition to the 27,600 ACs from Yan *et al.* ^12^ in the integrated dataset (**Supplementary Fig. 3a-b**). Utilizing the collected data, the integrated analysis annotated 63 AC types, revealing consistent expression profiles of known marker genes (**Fig. 3a-b** and Methods). While a minimal batch effect in each cluster was observed across different sample sources, CD90.1^+^ and Ma *et al.* RGC samples showed biased enrichment towards GABAergic types except for AC4, AC10, and AC28 (**Supplementary Fig. 4e**). The bias in cell type population appears to be directly tied to the preferential expression of *Thy1* (CD90) in sub-populations of ACs (**Supplementary Fig. 4d**). In particular, *Thy1* is characterized as being expressed primarily in GABAergic AC types ^26^.

**Figure 3.**
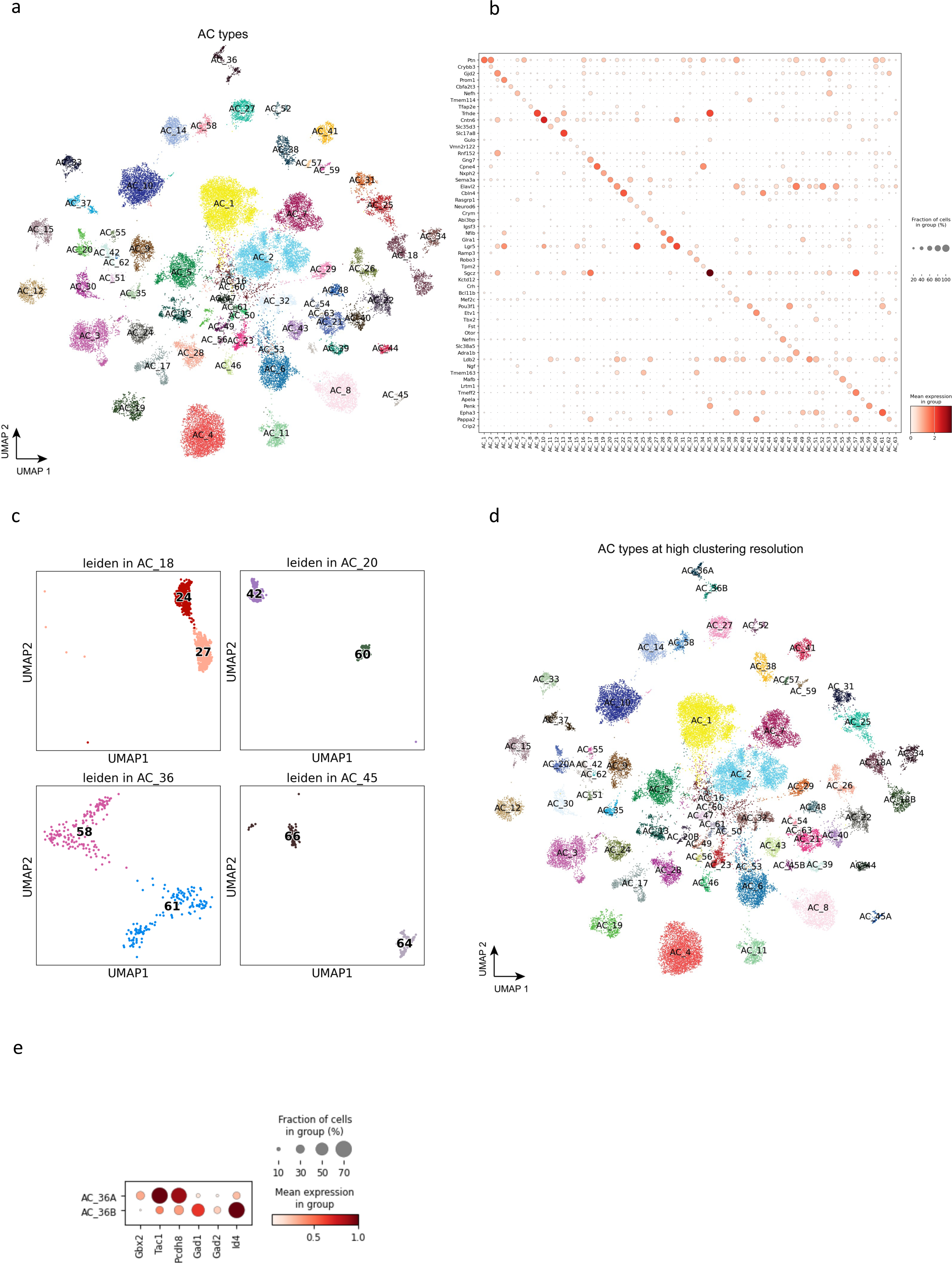
Amacrine cells. **(a)** UMAP visualization of AC cells colored by the annotated types. **(b)** Dot plot of canonical marker gene expression in AC types. **(c)** Four previously under-clustered AC types, i.e., AC18, AC20, AC36, and AC45, are split into two distinct clusters at a high resolution of clustering. **(d)** Visualization of AC cells colored by AC types at a high clustering resolution. **(e)** Dot plot of DEGs expressed in two split clusters for AC_36, stratifying *Gbx2^+^* AC types in AC_36.

The integrated analysis of ACs demonstrated that four types, AC18, AC20, AC36, and AC45, have been previously under-clustered, each splitting into two clusters in the integrated UMAP (**Fig. 3c**). AC18, which expresses *Cck* neuropeptide ^27^, is split into C24 and C27 in our clustering and has been labeled as AC18A and AC18B in the final annotation (**Fig. 3d**). Interestingly, the cell type marker *Cck* is highly expressed in AC18A, but not in AC18B (**Supplementary Fig. 5a**). AC20, which does not contain any known marker, is divided into C42 and C60 (AC20A and AC20B), with its marker *Sema3a* also expressed highly in AC20A, but not in AC20B (**Supplementary Fig. 5b**). A non-GABAergic non-glycinergic (nGnG) type 4, AC36, is split into C58 and C61 (AC36A and AC36B), consistent with previous finding of two morphologically distinct AC36 types in the INL and displaced in the GCL, stratifying to S3 and S5 sublaminae of the IPL ^10,28^. By examining the list of differentially expressed genes (DEG) between the two broadly isolated types ^28^, we annotated AC36A as the S3 type by the increased markers such as *Gbx2*, *Tac1*, and *Pcdh8* and AC36B as the S5 type by *Gad1*, *Gad2*, and *Id4*. (**Fig. 3e**). Lastly, a catecholaminergic type 1 cell type ^29^, AC45, is split into C64 and C66 (AC45A and AC45B). The expression of *Chl1*, which distinguishes catecholaminergic type 1 from type 2, was increased specifically in AC45A. The DEG analysis between the clusters of the previously under-clustered cell type revealed many genes enriched specifically in each cluster, with *Cck*, *Sema3a*, *Chl1* being one of the top-ranked genes in AC18A, AC20A and AC45A, respectively (**Supplementary Fig. 5b**). Out of the four under-clustered cell types, only one, AC20, showed a biased sample source from Yan *et al.* data. Furthermore, while cells from Yan *et al.* were distributed across both AC45A and AC45B, AC45B contains an increased number of cells from the newly generated CD90.1 sample (**Supplementary Fig. 4e**).

As a result, we have identified 67 AC types that can be grouped into four AC subclasses: 49 GABAergic, 10 Glycinergic, 3 Both, and 5 nGnG ACs. Within the final dataset, GABAergic ACs make up 67.7% of the total AC population, followed by Glycinergic ACs at 22.5%, GABA/Glycinergic ACs at 1%, and nGnG ACs at 8.7%. However, these distributions are likely biased towards GABAergic ACs due to the inclusion of cells from CD90.1^+^ and CD90.2^+^ enriched collections.

### Retinal ganglion cells

The integrated data contains 77,900 RGCs, primarily from the three publicly available datasets. The integration of the collected data identified all 46 previously identified RGC types (**Fig. 4a** and Methods). Examination of known cell type markers in the integrated data with the final annotation showed proper expression profiles in corresponding types ^9,18,30^ (**Fig. 4b**). Although no novel cluster was identified, our integrated analysis of RGCs similarly identified the division of two cell types, 16_ooDS_DV (ON-OFF direction-selective dorsal and ventral) and 18_Novel, into distinct clusters (**Fig. 4c**). The 16_ooDS_DV, which contains both types with dorsal and ventral orientation selective functional roles ^31,32^, was split into C31 and C39, similar to the supervised clustering analysis done in the Tran *et al.* ^9^, Jacobi *et al.* ^13^, and Ma *et al.* ^14^ studies. Examination of the marker genes *Calb1* and *Calb2* demonstrated that C39 is the ventral selective type with high expression of *Calb2*, and C31 is the dorsal selective type with *Calb1* expression ^9^. In addition, the 18_Novel type could also be split into C36 and C40. Interestingly, while C40 contained only cells with 18_Novel labels, C36 contained a mixture of 18_Novel and 44_Novel labels (**Supplementary Fig. 7a-c**). The same annotation improvements were also observed in Ma *et al.*^14^. Examination of 18_Novel markers *Pcdh20* and *4833424E24Rik* revealed increased expression of both markers in C40, yet *Pcdh20* expression was absent in C36 (**Supplementary Fig. 7e**). The DEG analysis further demonstrated many genes selectively expressed in these two clusters (**Supplementary Fig. 7d**). In total, we have identified 47 RGC types in the MRCA (**Fig. 4d**).

**Figure 4.**
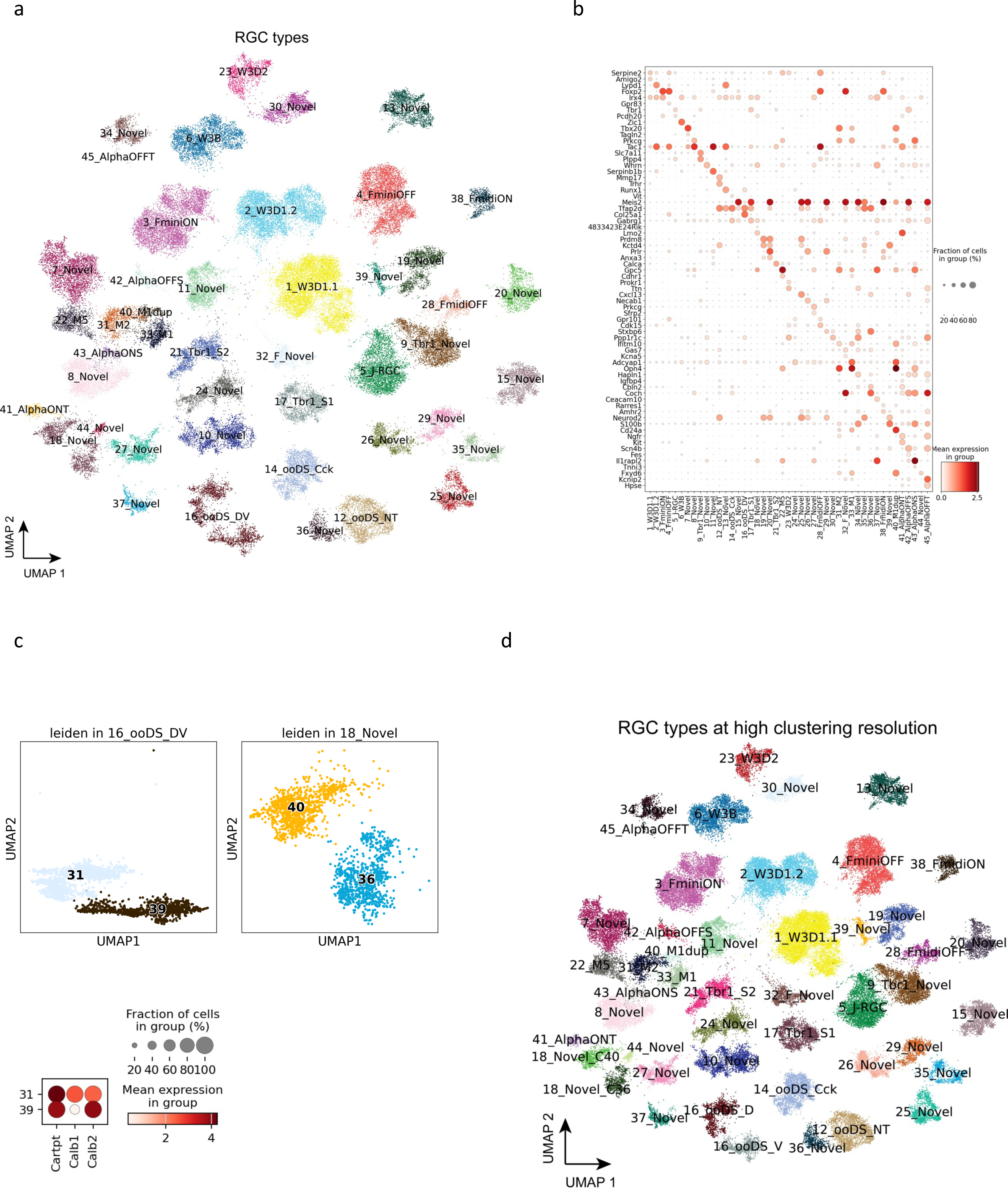
Retinal ganglion cells. **(a)** UMAP visualization of RGC cells colored by the annotated types. **(b)** Dot plot of canonical marker gene expression in RGC types. **(c)** Two previously under-clustered RGC types, i.e., 16_ooDS_DV and 18_Novel, are split into two distinct clusters at a high resolution of clustering. Dot plot of *Calb1* and *Calb2* in the two split clusters of 16_ooDS_DV. **(d)** Visualization of RGC cells colored by RGC types at a high clustering resolution.

### Non-neuronal retinal cells

To include the comprehensive set of cell types in the retina in the MRCA, 18,500 non-neuronal cells were integrated for six non-neuronal cell types, including astrocyte, endothelial, MG, microglia, pericyte, and RPE (**Supplementary Fig. 8a** and **Fig. 1e**). These cells are evenly distributed in the collected datasets, except for astrocytes solely from the Benhar *et al.* dataset^15^ (**Supplementary Fig. 8b**). After being combined with neuronal retinal cells, the MRCA consisted of 12 major classes and 138 cell types.

### Data dissemination at accessible interactive web browsers

The MRCA has been made available for public access using the CELLxGENE platform (https://cellxgene.cziscience.com/collections/a0c84e3f-a5ca-4481-b3a5-ccfda0a81ecc and https://mouseatlas.research.bcm.edu/) (**Fig. 5a-c**). The MRCA is also accessible on UCSC Cell Browser (https://retina.cells.ucsc.edu) and the Broad Single Cell Portal. Pre-computed gene expression profiles of all cells included in the integrated analysis can be examined and visualized. Users also have access to the metadata information, including major class and cell type labels in the database. The accessible interactive web browsers of the MRCA can aid in easy access to the transcriptome profiles of any given mouse retinal cells without the bioinformatic burden and provides a valuable tool for the vision community.

**Figure 5.**
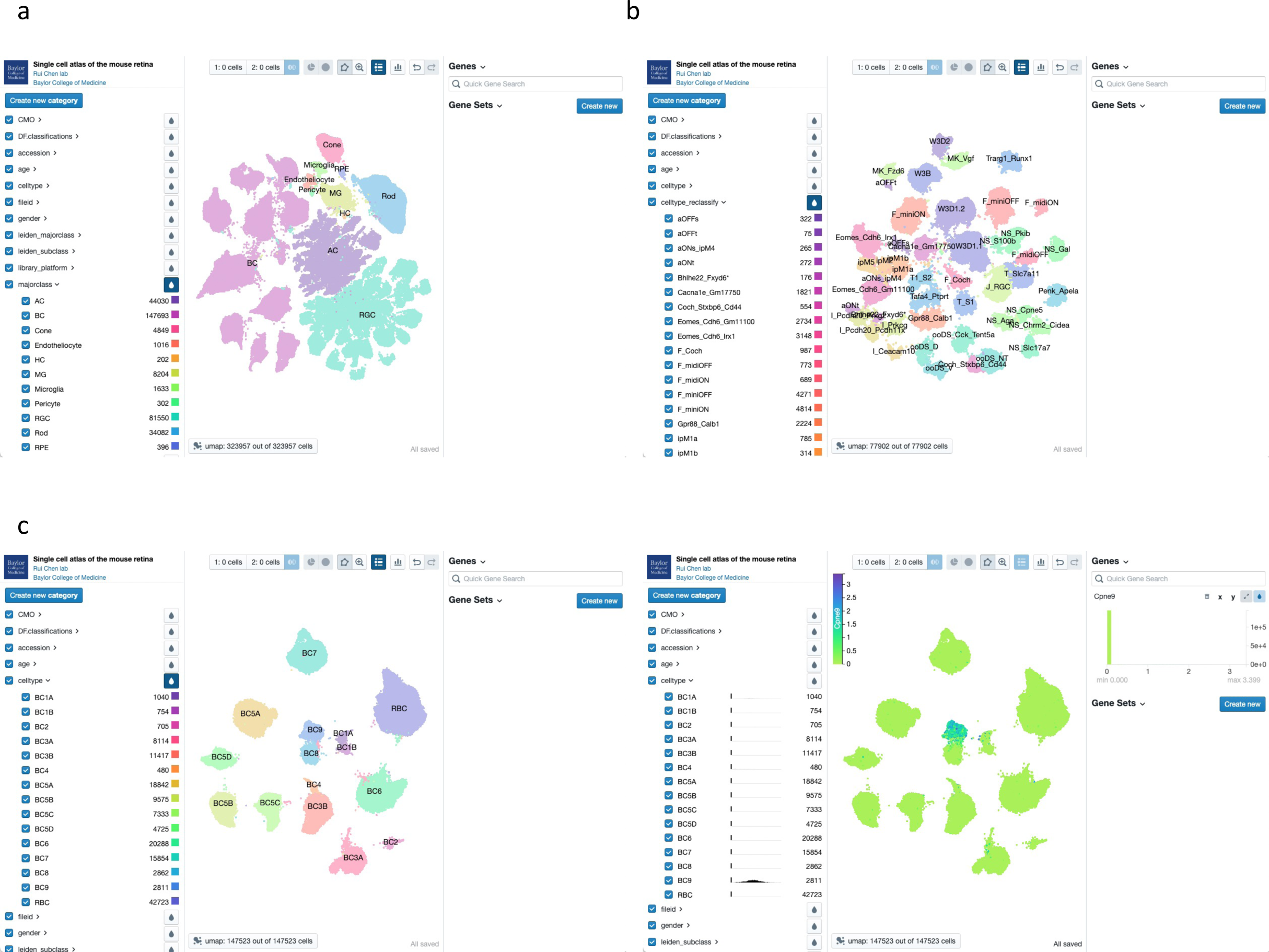
Visualization of MRCA in accessible interactive browsers. **(a)** Visualization of the MRCA in the CELLxGENE browser. The homepage depicts three panels to explore the MRCA. The left panel contains the pre-computed features facilitating the selection of cells by interested categories. The middle panel is the UMAP of the MRCA, colored by the annotated major classes. The right panel allows input of quick gene symbols and gene sets. **(b)** Visualization of the subclass RGC atlas in the CELLxGENE browser. The middle panel depicts RGCs colored by the reclassified names selected in the left panel. **(c)** Visualization of gene expression for a BC9 marker, *Cpne9*, in the BC atlas. The left subfigure shows the BC types, and the right subfigure highlights the normalized gene expression values of *Cpne9* for BC9 type in the middle panel.

## Discussion

As part of the central nervous system, the retina contains numerous neuronal types with distinct morphologies and functional roles ^1,33^. The heterogenous cell type composition and the stereotypically patterned structure of the tissue makes the retina an ideal model for single-cell sequencing studies in establishing the single-cell atlas ^7,22,34^. Although several scRNA-seq studies focusing on the retina tissue have been done previously ^7,9,11–15^, each available dataset contains single-cell profiles primarily of one or a few retinal cell classes with a limited number of cells. Furthermore, no systematic evaluation or comparison of the datasets has been done yet to cross-validate the cell type transcriptomes and address annotation consensus.

In this study, we generated scRNA-seq profiles of 189,000 retinal cells from 16 scRNA-seq experiments to perform an integrated analysis with 141,000 retinal cells from six previously reported datasets. Six out of the newly generated collections were done using endogenous retina tissues with simple dissociation and without enrichment. Photoreceptors constitute over 70% of the cell proportion in the retina ^2,35^, and there are only two subclasses of photoreceptors, which are well studied. Therefore, we utilized two methods for rare population cell type enrichment. The first way was depleting the rod photoreceptors. To achieve this goal, the rod photoreceptor cell surface marker, CD73, was used in seven of the 16 experiments. Though this marker is generally considered as a specific marker for rod photoreceptors, it is also expressed on the surface of a subset of ACs, HCs, and MGs. Depletion increased the enrichment of BCs from 12% to 90%. Furthermore, CD90.1 was used to enrich certain retinal neurons such as ACs and RGCs in three experiments. Enrichment of retinal cells with CD90.1 also showed an increased number of ACs with some RGCs.

One of the challenges in integrating and comparing publicly available data is that they are generated using different single-cell experimental platforms and analysis pipelines ^36,37^. One public data enriched with BCs from Shekhar *et al.* ^11^ was generated using the Drop-seq ^7^ technology and was processed separately using the Cell Ranger transcript annotation. The four other sources of publicly available data were done using the 10x Genomics platform. A minimal batch effect across data sources was observed in the integrated analysis, with the expected distribution and clustering of major classes from corresponding sources. While the newly generated data without enrichment were primarily composed of rod photoreceptors, cells from the newly generated data with enrichment and publicly available data showed a proper distribution across BCs, AC, and RGCs.

Integrated analysis of various scRNA-seq datasets allowed us to examine AC, BC, and RGC types, which together comprise over 100 distinct cell types. Through the integrated analysis, we addressed two key questions on the neuronal cell types in the retina: to confirm the consensus cell type signatures and to examine whether the total number of cell types of retinal neurons is exhausted. Following the initial integrated analysis to identify major classes, subsets of each major class were subjected to further integration and two-level clustering to annotate all previously identified cell types, which showed an even distribution of data sources in general. The cell type annotation was achieved through examining known marker gene expressions and previous annotation labels when available. Although our newly generated data resulted in a significantly increased number of cells in the integrated analysis of BCs, ACs, and RGCs, we did not observe significant increases of novel cluster. As such, the previously reported set of BC types in the adult mouse retina is likely complete, supported by the more than 7-fold increase in BCs in the integrated data. On the other hand, our integrated analysis updates annotations of AC and RGC types. In particular, we observed several instances of previously under-clustered AC and RGC types splitting into distinct clusters in our analysis. For example, we confirmed the separation of 16_ooDS_DV types into two distinct clusters in the integrated data of RGCs, which was separated into dorsal and ventral selective types only through supervised clustering in the Tran *et al.* study ^9^ and later confirmed in Jacobi *et al*. ^13^ study. Furthermore, we identified the separation of AC36 and assigned its clusters to S3 and S5, stratifying *Gbx2^+^* AC types^28^, which strengthens our analysis by connecting to biologically distinct cell types. The separation of previously merged cell types into distinct clusters can be attributed to the increased number of cells in our integrated analysis. This suggests that, while our AC and RGC type annotations are comprehensive, they will likely continue to be refined by future studies.

Finally, we have deposited the MRCA into interactive web browsers that are user-friendly and publicly accessible. This allows for the examination of raw and normalized gene expression profiles of all retinal cells, along with their metadata such as major class and cell type annotation. The MRCA not only provides the consensus signature of mouse retinal cell types by comparing multiple scRNA-seq data but also alleviates the bioinformatics burden for many vision researchers who wish to examine transcriptome signatures in any cell type of their interest.

## Methods

### Generation of scRNA-seq datasets of the mouse retina

We have generated 16 scRNA-seq samples of the mouse C57BL/6J retina (**Table 1**). All mice were male. All procedures were approved by the Institutional Animal Care and Use Committee (IACUC) and followed the Association for Research in Vision and Ophthalmology (ARVO) Statements for the Use of Animals in Ophthalmic and Vision Research, in addition to the guidelines for laboratory animal experiments (Institute of Laboratory Animal Resources, Public Health Service Policy on Humane Care and Use of Laboratory Animals). After dissection, retinas were dissociated into single cells using papain-based enzyme following the published protocol^38^. With activated 45U of papain (Worthington, Cat. #LS003126) solution (1mg L-Cystine, Sigma; 8 KU of DNase I, Affymetrix; in 5 ml DPBS), retina was incubated at 37C for ∼20min, followed by the replacement of buffer with 2ml ovomucoid solution (15 mg ovomucoid, Worthington Biochemical; 15 mg BSA Thermo Fisher Scientific; in 10 ml DPBS) and 500ul deactivated FBS. Following the enzymatic digestion step, the retina tissues were carefully triturated and filtered using 20 um plastic meshes. Trituration steps were repeated with additional 1ml ovomucoid solution until no tissue was visible. Single-cell suspension was spun down at 300g, 4C for 10 min and used in the next step.

To deplete the photoreceptors, cells were resuspended in 0.5% BSA and stained with CD73-PE antibody (MACS, Catalog: 130-102-616) for 10min at 4C (for each million cells, add 98ul 0.5% BSA with 2ul CD73-PE antibody) and washed with 35 ml 0.5% BSA at 4C for 10min. After being stained with Anti-PE microbeads (MACS, Catalog: 130-105-639) (80ul 0.5% BSA and 20ul microbeads per each million cells) for 15 min at 4C, cells were washed and resuspended in 0.5% BSA. CD73 negative neuronal cells were enriched by autoMACS Pro Separator (Miltenyi Biotec) DEPLETES mode. Similarly, CD90.1 positive neuronal cells were enriched with CD90.1 microbeads (MACS, LOT: 130-094-523; 90ul 0.5% BSA and 10ul CD90.1 microbeads per each million cells) and autoMACS POSSEL-S mode. Cells viability was 87%-94% when checked using DAPI staining under microscope.

Guided by 10X manufacturer’s protocols (https://www.10xgenomics.com), single-cell cDNA library was prepared and sequenced. Briefly, single-cell suspension was loaded on a Chromium controller to obtain single cell GEMS (Gel Beads-In-Emulsions) for the reaction. The library was prepared with Chromium Next GEM single cell 3’ kit V2 (10X Genomics) and sequenced on Illumina Novaseq 6000 (https://www.illumina.com). Our newly generated single cell data was sequenced at the Single Cell Genomics Core at Baylor College of Medicine.

### Data collection and preprocessing of the mouse retinal scRNA-seq

To recover high-quality cells, data samples were processed through a quality control pipeline (https://github.com/lijinbio/cellqc). In brief, raw sequencing reads of 10x Genomics were first analyzed by the 10x Genomics Cell Ranger pipeline (version 7.0.1) ^39^ using the mm10 genome reference obtained from 10x Genomics (https://cf.10xgenomics.com/supp/cell-exp/refdata-gex-mm10-2020-A.tar.gz). Potential empty droplets in the filtered feature count matrices were further detected by dropkick ^40^. Background transcripts contamination in the retained true cells were eliminated using SoupX ^41^. DoubletFinder then was utilized to estimate and exclude potential doublets with high proportions of simulated artificial doublets ^42^. In the resulting singlets, we extracted high-feature cells that contain ≥ 300 features, ≥ 500 transcript counts, and ≤10% of reads mapped to mitochondrial genes.

In addition to our own data, we have incorporated well-characterized public datasets. Specifically, we have integrated cell-type-enhanced profiling data for amacrine cells (accession: GSE149715) ^12^, bipolar cells (accession: GSE81904) ^11^, and retinal ganglion cells (accession: GSE133382) ^9^. Furthermore, we have included four samples from wild-type mice were also collected from GSE201254 to account for retinal ganglion cells ^13^. To account for non-neuronal retinal cells, nine control samples were collected from GSE199317 ^15^. These cell-type specific single-cell datasets form the basis for subclass clustering in our mouse retina reference. To generate the updated transcriptome measurement of the GSE81904 from Shekhar et al., which was derived from the Drop-seq protocol, we applied the Drop-seq pipeline using the source code available at https://github.com/broadinstitute/Drop-seq. To ensure consistent gene feature annotation with the Cell Ranger pipeline, we used the gene annotation GTF file from the 10x Genomics mm10 genome reference package during the alignment of Drop-seq reads. In addition, GSE149715, GSE133382, GSE201254, and GSE199317 were also processed from scratch using raw sequencing reads using the 10x Genomics Cell Ranger pipeline (version 7.0.1) ^39^. To incorporate the high-quality cell type annotation of four public datasets, released count matrices and cell labeling were downloaded for meta-analysis. To further eliminate potential multiples in the integrated analysis, Solo doublet detection algorithm was used to identify potential multiples.

### Data integration of scRNA-seq datasets

To eliminate technical variations in samples derived from different studies and experiments, 52 samples were integrated to remove the batch effect by scVI ^43^. scVI explicitly formulates the batch effect as a latent variable in the deep generative model of observed expressions. Normalized expression was applied to detect highly variable genes (HVGs) using the Seurat algorithm (flavor: seurat). The “sampleID” was used as the batch key for calculating HVGs and the batch variable in the scVI modeling. The scVI model utilized 2 hidden layers (n_layers: 2) and a 30-dimensional latent space (n_latent: 30). The trained low-dimensional representation was used for cluster detection with the Leiden algorithm ^44^. UMAP of low-dimensional visualization was generated by the Scanpy package ^45^.

### Cell clustering and cell type annotation

To annotate major classes of cell clusters, we incorporated well-annotated cell labels released from public datasets, i.e., Yan *et al.* for ACs, Shekhar *et al.* for BCs, and Tran *et al.* and Jacobi *et al.* for RGCs. Cells from Yan *et al.* were annotated into 63 AC types. Cells from Shekhar *et al.* were 15 BC types showing in 14 clusters with small numbers of cells annotated as ACs, rod, and cone. Tran *et al.* cells were identified as 45 RGC types. The cell type labels of these well-annotated cells are used to annotate integrated cell clusters. To annotate isolated cell clusters that were isolated from existing cell labels of the public datasets, cluster-specific markers were examined from the top ranked genes generated by the Wilcoxon rank-sum test using the rank_genes_groups() function in the Scanpy package ^45^.

To annotate subclass BC, AC, and RGC, subclass-specific cells were isolated and integrated using scVI. The generated low-dimensional embeddings were used to detect clusters using the Leiden algorithm. To determine the optimal number of clusters for subclasses, a two-level clustering approach was applied. In the first level of clustering, various resolutions were tested to achieve clustering without over-clustering in UMAP visualization. The second-level clustering refines the clusters from the first-level clusters by testing various resolutions to achieve optimal clustering without over-clustering on UMAP again. In the first-level, Leiden clusters containing the majority of one type were annotated. When Leiden clusters contained more than one types, cells within the clusters were isolated. Within each subset of isolated cells, Leiden clusters were calculated again using the same low-dimensional embedding. The second-level Leiden clusters were examined for their cell label to determine their cell types.

To construct the BC atlas, data samples for BCs were integrated using scVI. Initially, 33 clusters were identified, of which 30 could be matched and merged to individual BC types by examining previously generated cell labels and their known marker gene expression ^11,24^, while the remaining 3 clusters (C30, C31, and C32) were excluded from the analysis as they contained non-BCs from previous annotation labels or had high UMI counts (**Fig. 2a** and **Supplementary Fig. 2a-c**). Consequently, 15 BC types were identified and annotated.

To construct the AC atlas, the data integration analysis for ACs using scVI identified a total of 71 clusters, of which 62 clusters could be matched and merged to 49 individual AC types via previous annotation labels and known marker expression. However, 8 clusters were over-clustered that contained two or more previous AC type labels, and one cluster (C70) was excluded from the AC reference due to non-AC cells (**Supplementary Fig. 3c-d**). To further address the 8 remaining over-clustered clusters (**Supplementary Fig. 4a**), we utilized a two-level annotation approach. This involved isolating cells from each cluster and refining the clustering. The two-level annotation allowed the separation of the remaining 14 types: AC11, AC16, AC29, AC42, AC47, AC50, AC53, AC54, AC55, AC56, AC60, AC61, AC62, and AC63 (**Supplementary Fig. 4a-c**). This revealed clusters that primarily consisted of RGCs, which have been removed in the integrated AC map (**Supplementary Fig. 4c**). As a result, 63 AC types were identified and annotated.

Three AC types, AC16, AC53, and AC62, were identified as dual types expressing both canonical GABAergic and glycinergic receptors in the study by Yan *et al*. AC16, however, was shown as a suspected doublet in their study, alongside AC60. Similarly, our UMAP showed loose cluster formation of AC16 and AC60 in proximity to each other, with relatively high UMI counts (**Fig. 3a** and **Supplementary Fig. 4e**). In addition, our integrated UMAP showed AC53 cells spread out in the middle of AC6 cells. Although the AC53 cluster was resolved in the second-level annotation, the loose clustering of AC53 cells is quite apparent. The third dual type, AC62, was also under-clustered and merged with AC42 and AC55. While AC62 was resolved in the second-level annotation, AC62 also appears near its neighboring cluster, AC42, in the UMAP. With very few cells being annotated as dual types in CD90.1 and Ma *et al.* samples, which express high levels of *Thy1* (data not shown), further validations of the dual types are required.

To construct the RGC atlas in the MRCA, the integrated analysis identified 54 clusters with an even distribution of cells from different data sources in most clusters (**Supplementary Fig. 6a-d**). Out of these clusters, 48 can be mapped and merged into 39 individual RGC types previously identified using marker gene expression and previous annotation labels (**Supplementary Fig. 6a-b**), while five clusters were over-clustered that contained multiple previous RGC types, and one cluster (C8) contained a mixture of several RGC type labels with high UMIs and was excluded from the downstream analysis as multiplets. To annotate the remaining seven types found in the five clusters with multiple labels, the second-level annotation was performed, which resulted in a clear separation of all 46 previously identified RGC types (**Fig. 4a** and **Supplementary Fig.7a-c**).

### Differentially expressed gene analysis

To identify genes that are differentially expressed between cell types, we generated pseudo-bulk transcriptome of each annotated cell type in individual sample id. We used pyDESEQ2 ^46^ to compare two clusters or types using the Wald test and identified genes specifically expressed in each cluster or type. Differentially expressed genes are identified under *q-*value < 0.05. The Wald statistics (log2FoldChange divided by lfcSE) was used to rank and select the top 10 genes expressed in each type.

## Data Availability

The raw sequencing reads of sixteen newly generated samples have been deposited at NCBI GEO under the accession GSE243413. The landing page for the MRCA data resources is accessible at https://rchenlab.github.io/resources/mouse-atlas.html. Processed cell-by-gene count matrices, along with cell type annotations, are available on Zenodo. Furthermore, both raw and normalized count matrices and cell type annotations are publicly accessible on the CELLxGENE data collection at https://cellxgene.cziscience.com/collections/a0c84e3f-a5ca-4481-b3a5-ccfda0a81ecc. The MRCA is also hosted on the Baylor College of Medicine data portal at https://mouseatlas.research.bcm.edu. Additionally, access to the MRCA is provided on the UCSC Cell Browser at https://retina.cells.ucsc.edu and the Broad Single Cell Portal.

## Code Availability

All code used for the MRCA project can be found in the MRCA reproducibility GitHub repository (https://github.com/RCHENLAB/MouseRetinaAtlas_manuscript). The pipeline to process the unpublished and collected public datasets is accessible at https://github.com/lijinbio/cellqc.

## Acknowledgements

We thank Alice Tian for her meticulous proofreading of the manuscript. This project was funded by NIH/NEI R01EY022356, R01EY018571, S10OD032189, Chan Zuckerberg Initiative (CZI) award CZF2019-002425, RRF to R.C.

## Author contributions

J.L., J.C., X.C., and R.C. conceptualized and designed the study. R.C. supervised the study. X.C. and Y.L. generated scRNA-seq data in this study. J.L., J.M., and S.P. compiled dataset collection. J.L., J.C. and S.P. developed the integrated analysis pipeline and performed the integration and annotation analysis. J.R.S, G.M., and B.J.F. provided public datasets before publishing. N.M.T. provided input for various annotation. All authors wrote, reviewed, and contributed to the manuscript.

## Competing interests

The authors declare no competing interests.

## Notes

### Competing Interest Statement

The authors have declared no competing interest.

