## Supplementary Figures for "Comprehensive single-cell atlas of the mouse retina"

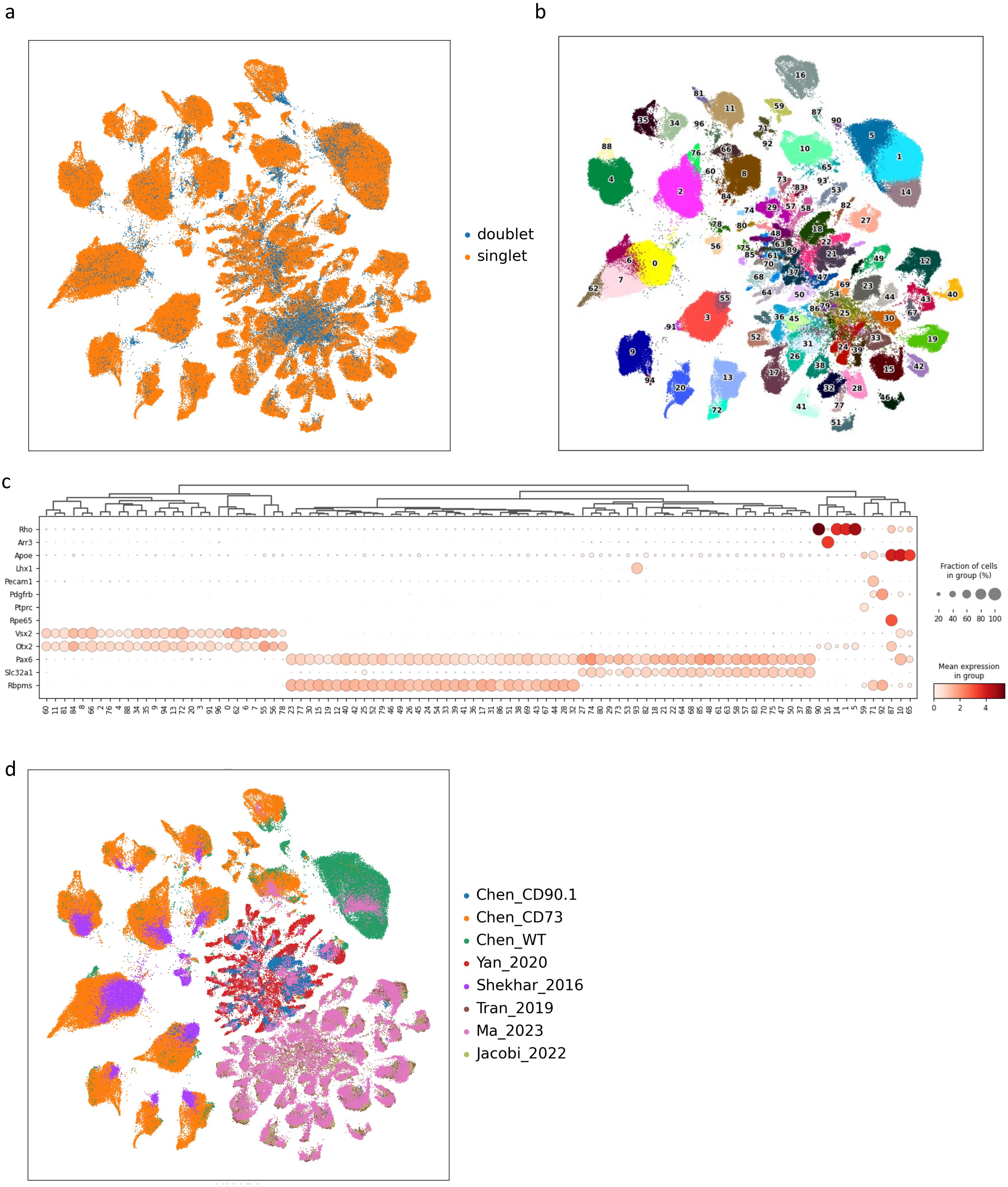


**Supplementary Figure 1. Cell type annotation of major classes**

**(a)** UMAP plot of the integrated data colored by doublet status using SOLO algorithm. Doublets are labeled in blue, while singlets are labeled in orange. (**b**) UMAP plot of the integrated data colored by each Leiden cluster. The Leiden cluster ID is placed on top of each cell cluster. **(c)** Dot plot of expression patterns of canonical markers of major classes in each Leiden cluster. Marker gene expression allows the annotation of 11 major classes. **(d)** UMAP plot of the integrated data colored by data sources. An even distribution of cells across clusters is shown by the expected data source.

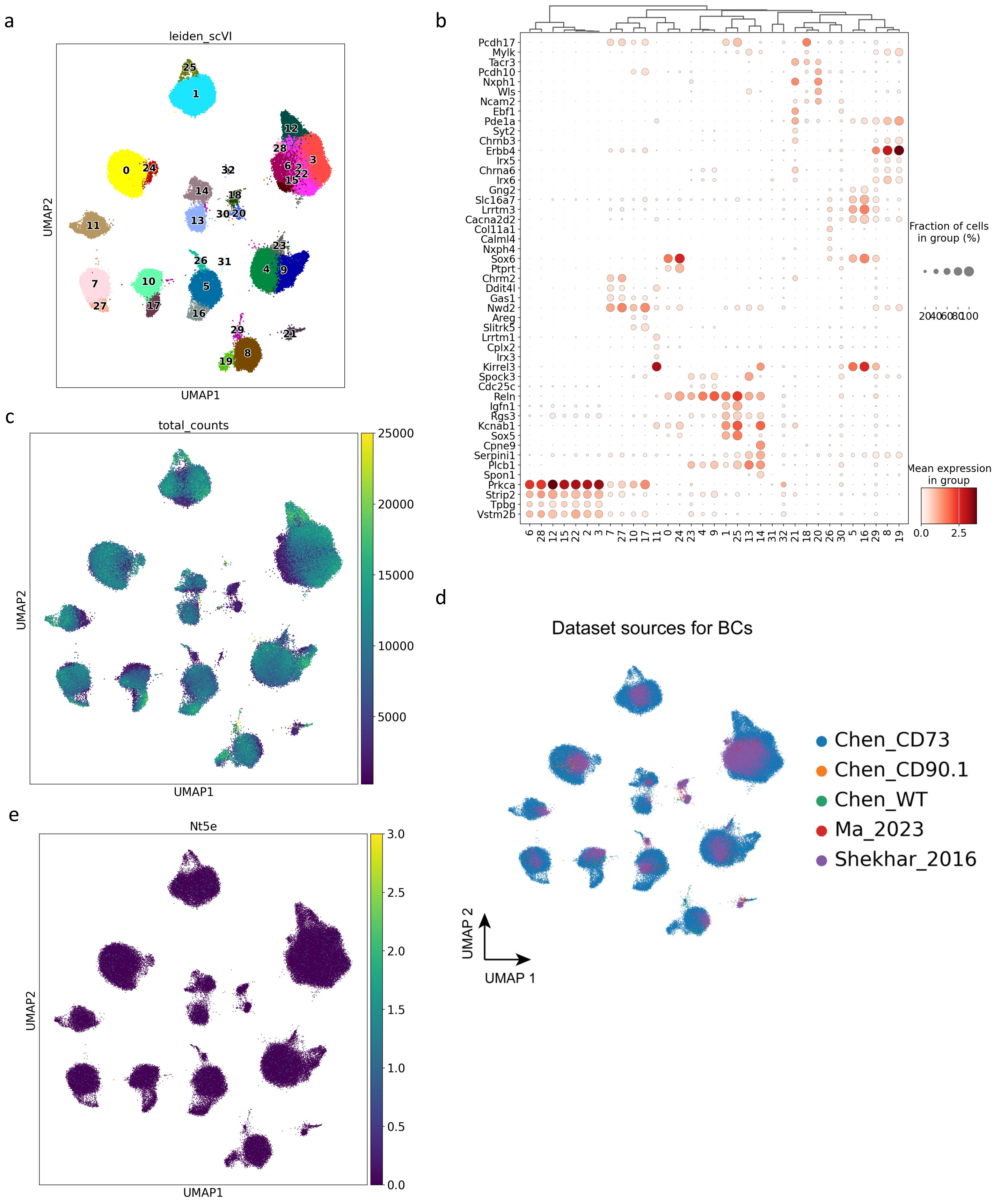


**Supplementary Figure 2. Cell type annotation of bipolar cells**

**(a)** UMAP visualization of BCs colored by cell clusters. **(b)** Dot plot of BC type marker gene expression in each cluster. **(c)** UMAP plot of BCs colored by the total UMI counts. **(d)** UMAP plot of BCs colored by data sources. **(e)** UMAP plot of *Nt5e* (CD73) expression in BCs.

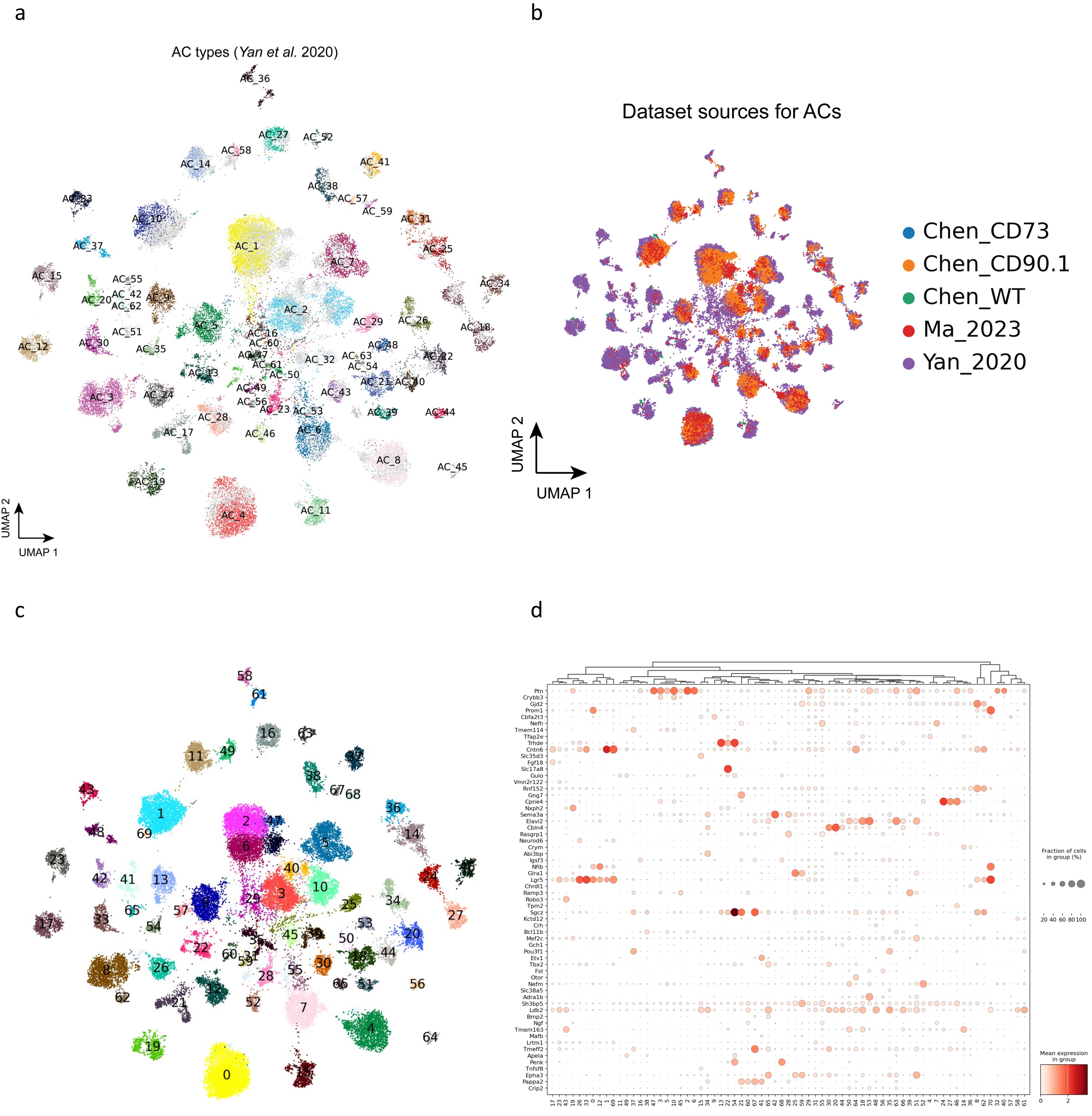


**Supplementary Figure 3. Cell type annotation of amacrine cells**

**(a)** UMAP visualization of ACs colored by public cell type labels from *Yan et al.* 2020. The newly discovered cells without public labeling are colored in gray. **(b)** UMAP plot of ACs colored by data sources. **(c)** UMAP visualization of ACs colored by 71 cell clusters. **(d)** Dot plot of AC type marker gene expression in 71 clusters.

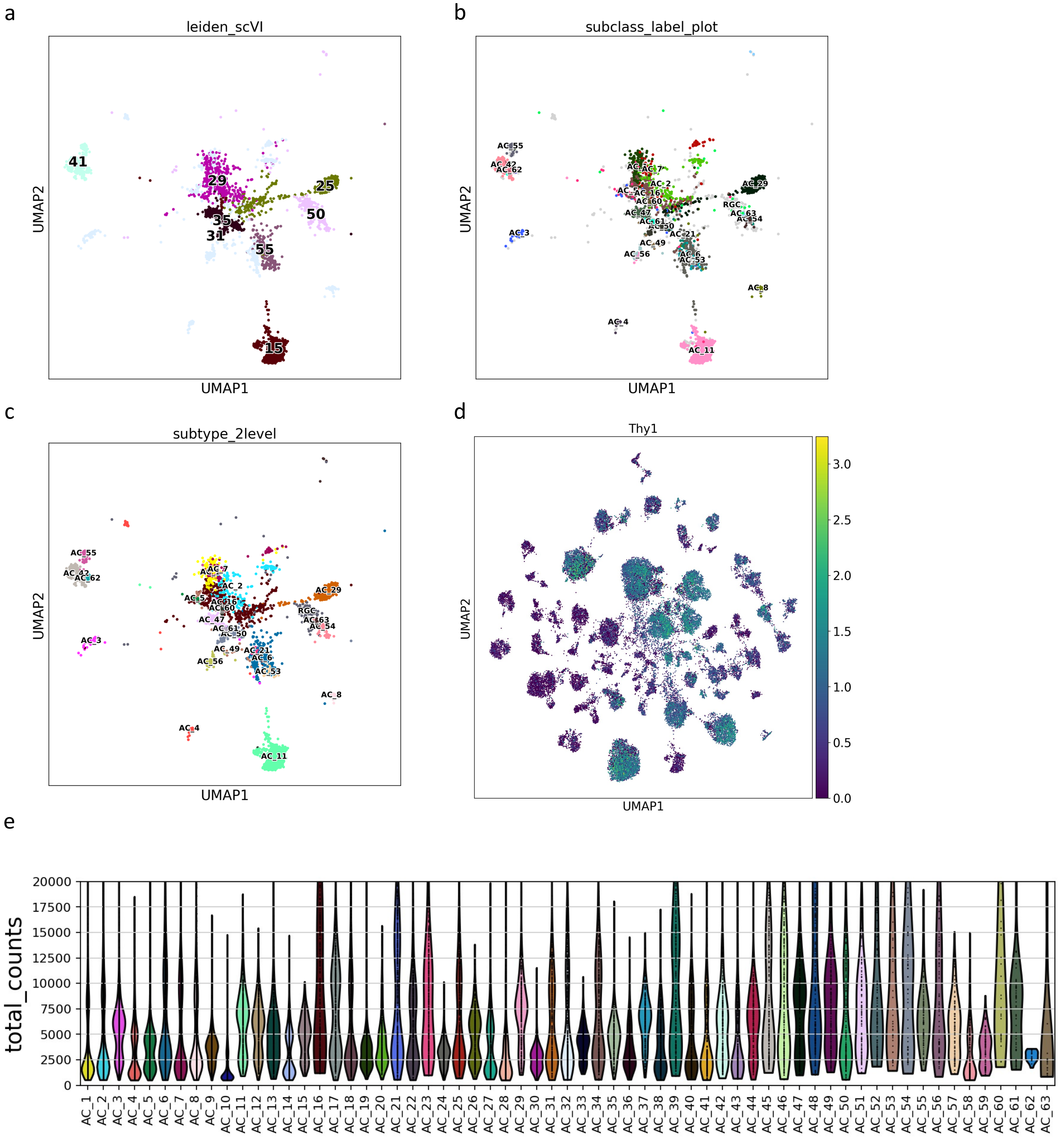


**Supplementary Figure 4. Annotation of over-clustered amacrine cells**

**(a)** UMAP visualization of the 8 cell clusters in ACs that contain more than one type, based on *Yan et al.* 2020. The 8 clusters are C15, C25, C29, C31, C35, C41, C50, C55. **(b)** UMAP visualization of the 8 clusters colored by public cell type labels from *Yan et al.* 2020. **(c)** UMAP visualization of the 8 clusters colored by AC types using the two-level annotation approach in this study. **(d)** UMAP plot of *Thy1* (CD90) expression in ACs. **(e)** Violin plot showing the UMI counts of the annotated AC types.

a

b

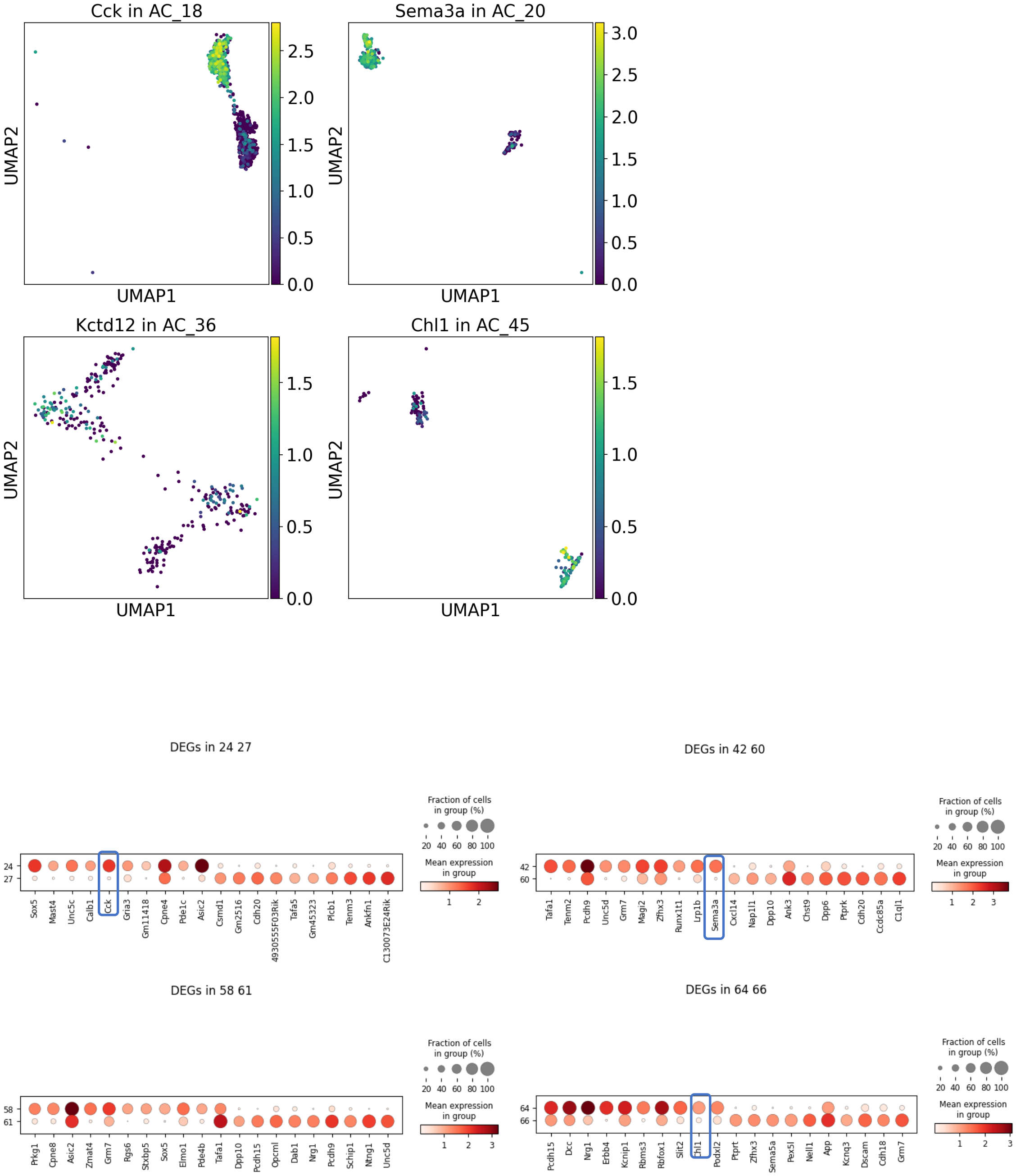


**Supplementary Figure 5. Annotation of under-clustered amacrine cells**

**(a)** Feature plot showing the marker gene expression of previously under-clustered AC types. **(b)** Dot plot showing the top 10 differentially expressed genes between split clusters for previously under-clustered AC types.

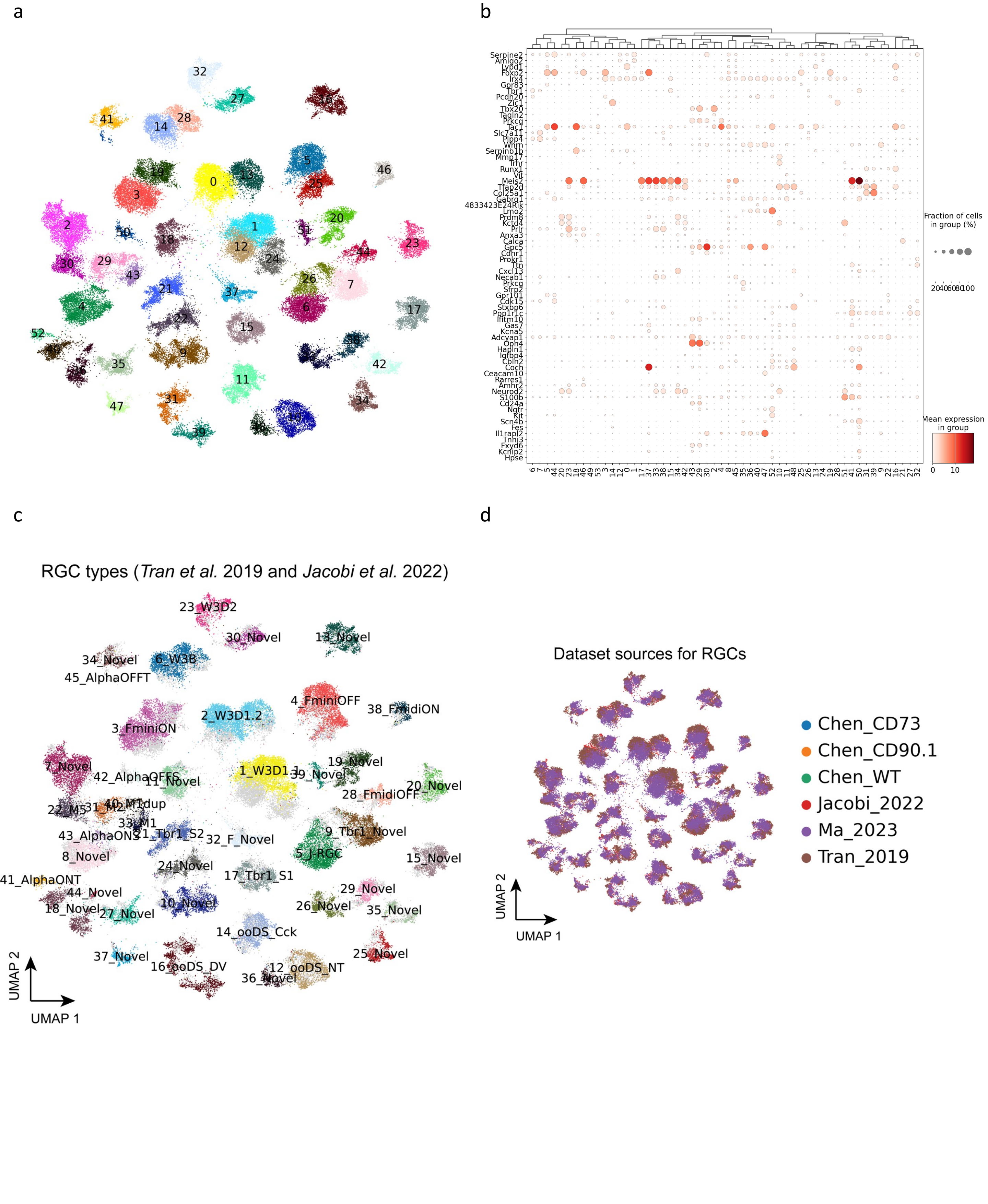


**Supplementary Figure 6. Cell type annotation of retinal ganglion cells**

**(a)** UMAP visualization of RGCs colored by cluster numbers. **(b)** Dot plot of RGC type marker gene expression in each cluster. **(c)** UMAP visualization of RGCs colored by public cell type labels from *Tran et al.* 2019 and *Jacobi et al.* 2022. The newly discovered cells without public labeling are colored in gray. **(d)** UMAP plot of RGCs colored by data sources.

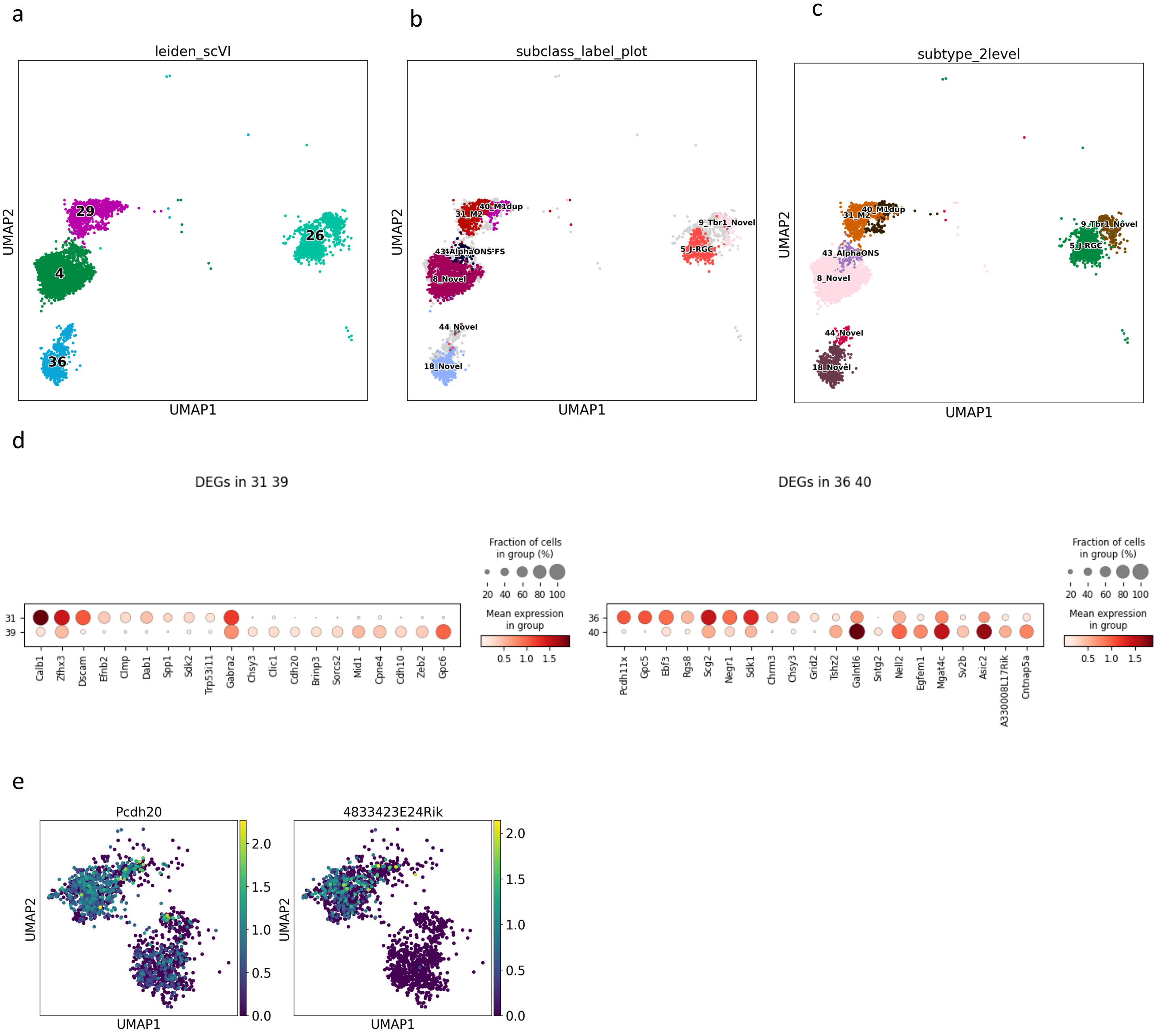


**Supplementary Figure 7. Annotation of over- and under-clustered retinal ganglion cells**
**(a)** UMAP plot of the 4 clusters in RGCs that contain more than one type, based on *Tran et al.* 2019 and *Jacobi et al.* 2022. **(b)** UMAP plot of the 4 clusters colored by public cell type labels from *Tran et al.* 2019 and *Jacobi et al.* 2022. **(c)** UMAP visualization of the 4 clusters colored by RGC types using the two-level annotation approach in this study. **(d)** Dot plot showing the top 10 differentially expressed genes between split clusters for previously under-clustered RGC types. **(e)** Feature plot showing the marker gene expression of 18_Novel type. Both markers are enriched in one cluster, but not the other.

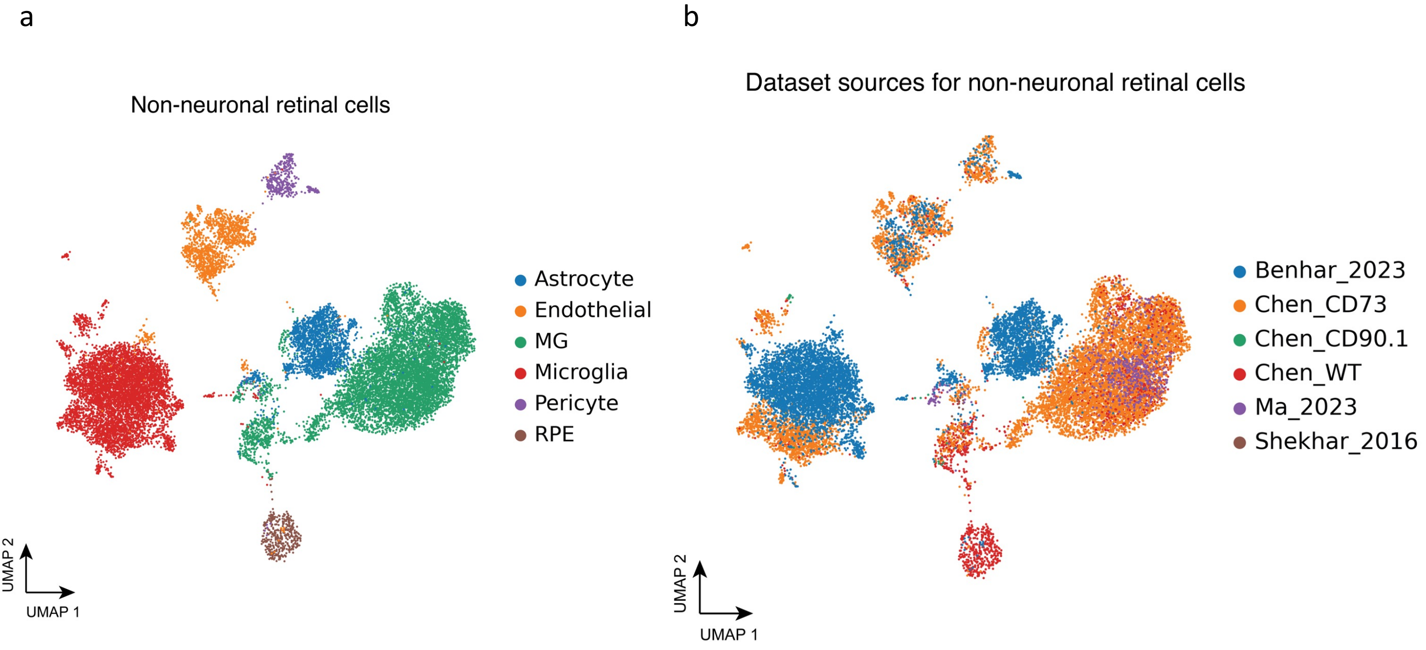


**Supplementary Figure 8. Non-neuronal retinal cells**

**(a)** UMAP visualization of non-neuronal retinal cells colored by major classes. **(b)** UMAP plot of non-neuronal retinal cells colored by dataset sources.
